# Clinical Forecasting using *Ex Vivo* Drug Sensitivity Profiling of Acute Myeloid Leukemia

**DOI:** 10.1101/2022.10.11.509866

**Authors:** Aram N. Andersen, Andrea M. Brodersen, Pilar Ayuda-Durán, Laure Piechaczyk, Dagim Shiferaw Tadele, Lizet Baken, Julia Fredriksen, Mia Stoksflod, Andrea Lenartova, Yngvar Fløisand, Jorrit M. Enserink

**Author notes:** Equal contribution (Aram N. Andersen, Andrea M. Brodersen, Pilar Ayuda-Durán).

## Abstract

Acute Myeloid Leukemia (AML) is a heterogeneous malignancy involving the clonal expansion of myeloid stem and progenitor cells in the bone marrow and peripheral blood. Most AML patients eligible for potentially curative treatment receive intensive chemotherapy. Risk stratification is used to optimize treatment intensity and transplant strategy, and is mainly based on cytogenetic screening for structural chromosomal alterations and targeted sequencing of a selection of common mutations. However, the forecasting accuracy of treatment response remains modest. Recently, *ex vivo* drug screening has gained traction for its potential in personalized treatment selection, as well as a tool for identifying and mapping patient groups based on relevant cancer dependencies. We systematically evaluated the use of drug sensitivity profiling for predicting patient survival and clinical response to chemotherapy in a cohort of AML patients. We compared computational methodologies for scoring drug efficacy and characterized tools to counter noise and batch-related confounders pervasive in high-throughput drug testing. We show that *ex vivo* drug sensitivity profiling is a robust and versatile approach to patient prognostics that comprehensively maps functional signatures of treatment response and disease progression. In conclusion, *ex vivo* drug profiling can accurately assess risk of individual AML patients and may guide clinical decision-making.

## Introduction

Acute Myeloid Leukemia (AML) is a heterogeneous cancer where the clonal expansion of myeloid progenitor cells (blasts) in the bone marrow and peripheral blood, interfere with healthy hematopoiesis resulting in immunodeficiency, thrombocytopenia and anemia^1^. The current 5-year survival rate of patients over 60 years of age is estimated to be 10-15%^2^. Most treatment-eligible individuals receive standard induction chemotherapy, which consists of a three-day treatment of 60 mg/m^2^ Daunorubicin (or Idarubicin) and 100-200 mg/m^2^ Cytarabine (or Ara-C) intravenously for seven days^3^. Survival rates of older patients have not substantially improved over the past decades, underscoring the need for better clinical assessment and more accurate prognostic approaches.

Current risk stratification methods such as the European LeukemiaNet (ELN) guidelines for AML stratification are mainly based on parameters such as molecular pathology and genomic alterations^4^. However, a substantial number of patients remains difficult to stratify, such as cytogenetically normal AML, and approximately 50% of patients are stratified as intermediate-risk patients for whom selection of the appropriate treatment regimen remains a major challenge^4–6^. Various non-genomic methods have been developed for identification of clinically and biologically relevant molecular subtypes for risk stratification, such as flow and mass cytometry, transcriptomics and proteomics^7–12^, although clinical implementation of these methods has generally been slow.

Recently, *ex vivo* drug sensitivity profiling has been used as a precision medicine approach to identify potential compounds that may be repurposed for the treatment of various types of cancer, including AML^13–19^. Here, leukemic cells derived from bone marrow aspirates or peripheral blood are incubated in the presence of various drugs at different concentrations. Drug responses are then quantified and overall drug efficacy is scored from a dose-response relation. This approach allows for rapid screening of a high number of compounds and has identified potential novel treatment avenues for AML.

The validity of drug sensitivity profiling requires that clinically relevant characteristics are conserved in *ex vivo* analyses; i.e. drug responses should reflect cancer dependencies, and drug sensitivities should be measurable reliably despite potential technical confounders and noise. Several quality control practices have been developed to ensure the fidelity of high-throughput drug testing^20–22^. A common procedure is to use model curve-fitting to de-noise dose-response data and summarize the drug sensitivities with the half-maximal effective concentration (EC50), or the area under the curve (AUC)^23,24^. Development of dose-range standardized function integrals based on the Hill equation, such as the drug sensitivity score (DSS), have improved the replicability of drug sensitivity profiles across independent studies^25–27^. However, curve-fitting of incomplete or non-sigmoidal drug responses remains a challenge for large-scale drug testing, and single output metrics from Hill models tend to yield incomplete information about the biological variability of a given drug response^23,28^. Therefore, streamlined analysis pipelines typically offer multiple alternative dose-response evaluation and error-reporting tools^22,29^.

While *ex vivo* drug screening of cancer cells using clinically relevant chemotherapeutics has been shown to predict treatment outcome, a systematic evaluation of the methodology and clinical use of drug sensitivity profiling in more expansive drug sets has not been done^30–33^. In this study, we evaluated the clinical information available in *ex vivo* drug sensitivity profiles within a cohort of 70 AML patients by using several statistical techniques and machine learning routines. Regularized regression was used to evaluate the clinical forecasting potential and risk-interpretability from *ex vivo* drug sensitivity profiles. We show that drug sensitivity profiling can improve patient risk stratification to aid clinical decision-making.

## Materials and Methods

### Patient cohort

Bone marrow aspirates and/or peripheral blood samples were collected from 70 adult patients diagnosed with AML and treated at the Department of Hematology at Oslo University Hospital in Norway between 2014-2017. Patients were included consecutively in this observational study. The study was performed in accordance with the Declaration of Helsinki and samples were collected following written informed consent. The study was approved by the Regional Committee for Medical Research Ethics South-East Norway (REK 2015/2012). Patient data that were collected included survival, sex, age, WHO patient performance status, routine diagnostic workup consisting of flow cytometry and genetic biomarkers investigated by G-banding, RT-qPCR, FISH and fragment analysis of FLT3 and NPM1, as well as ELN2022 risk stratification and FAB classification (see Suppl. Table S1 for a complete overview of patient characteristics). The median age was 60 years. All patients were subjected to standard treatment, which consisted of a 30 min infusion of anthracycline [daunorubicin (60 mg/m^2^) or idarubicin (10-12 mg/m^2^)] for three days in combination with a 24-hour infusion of cytarabine (Ara-C) for seven days, although the length and dosage of each treatment were in some cases altered depending on the patient’s age and general condition. Some patients received other drugs in addition to standard treatment (see Table S1 for details).

#### *Ex vivo* drug screens

*Ex vivo* drug sensitivity screens were performed on freshly isolated blast cells from bone marrow- or peripheral blood from 70 patients as previously described^8^. See Suppl. Methods for details.

### Dose-response analysis

Dose response metrics and quality control measures were calculated using R and Breeze^22^. See Suppl. Methods for details.

### Clinical data processing

The clinical data were grouped into two feature sets, with binarized dummy variables for categorical data. Clinical prognostic features included age at the time of diagnosis, sex, WHO patient performance status, and ELN2022 risk stratifications. Genetic features included specific mutations and chromosomal rearrangements with coverage for at least three patients. Missing data were zero-imputed. The survival times were computed from the date of diagnosis until the registered date of death or the last recorded visiting date. Response to induction was binarized based on the persistent presence of AML or reaching complete remission after induction therapy. Relapse was binarized for the sub-population of patients having reached complete remission.

### PCA and RCPC

PCA was performed using singular value decomposition (SVD) on scaled matrices of drug sensitivity metrics (or z-scores), and RCPC was performed by reconstructing the datasets by removing the leading singular values and singular vectors. The association of specific sample characteristics with a principal component was measured from the adjusted r-squared using linear regression, and the cumulative variance explained was computed as the cumulative sum of adjusted r-squared values multiplied with the proportion of variance explained of their respective principal components. See Suppl. Methods for details.

### Model training and cross-validation

Regularized Cox models were trained on the different drug sensitivity metrics and the binarized clinical variables using the glmnet package^48^. Ridge, Lasso, or Elastic net models were trained by setting the penalty mixture parameter (⍰?? to 0, 1, or 0.4 respectively, unless otherwise specified, and screening over a sequence of penalties using leave-one-out cross-validation (due to the low sample size). The model with the best average cross-validation deviance score was selected for testing or further analysis of model coefficients. The Cox model coefficients represent the change in log-hazard ratio (log-HR) as a function of drug sensitivity. Regularized binomial models were used for logistic regression of binarized clinical outcomes, and optimized using the same procedure.

Pre-selection of variables prior to model training was based on thresholds for standard deviations in drug sensitivity (as indicated), where the number of drugs with the greatest drug sensitivity standard deviations were selected. For a random sampling of features (100), the rAUC standard deviation was used as sampling weights.

### Model testing

For testing the models, a random sample of 10 patients was withheld from the training procedure, keeping a 4/10 proportion of deceased patients to maintain the approximate proportions in the original dataset. The prediction accuracy on test data was scored with Harrell’s concordance index (C-index), and the training and testing procedure was repeated 50 or 200 times, as indicated^49^. For testing of models predicting binary clinical outcomes, a five-fold training and testing procedure was performed and prediction accuracy was measured using an area under the receiver operating characteristic curve (ROC AUC).

### Variable importance

The importance of individual drugs was assessed by their contribution to predictivity through a variable withdrawal test, and the statistical significance of the model coefficients. The contribution to predictivity was measured by training and testing models leaving out individual drugs one-by-one and computing the change in C-index from the model containing all drugs. The statistical significance of model coefficients was estimated by training 200 models on bootstrapped datasets and computing the mean and confidence interval for the fitted coefficients.

### Model coefficient clustering

Estimated Ridge coefficients from models predicting different clinical outcomes were standardized and drugs with a coefficient of at least three standard deviations from the mean were selected for hierarchical clustering using euclidean distance and ward.D2.

### Enrichment test

Parametric enrichment analysis was performed using the GSEA function from ClusterProfiler based on ranked Ridge model coefficients or differential drug sensitivity z-scores, where drug sets were defined based on drug target or class association^50^.

### Drug sensitivity clustering and patient risk stratification

Hierarchical clustering was performed on a full drug sensitivity dataset, or the same data weighted by the respective Lasso coefficients for each drug, resulting in a dimension-reduced (by removing zero-sum rows) survival association matrix. Clustering was performed using euclidean distance and ward.D. For unsupervised classification the dendrograms were divided into 8 groups for the drug dimension and 6 groups for the patient dimension. Correspondence with drug annotations or patient annotations was measured using the Rand index. For patient annotation based on mutation profiles, a separate unsupervised classification using the same hierarchical clustering procedure of the binary mutation matrix was performed.

## Data and code availability

The data used in this study are available upon reasonable request. All code is freely available at https://github.com/Enserink-lab/DSCoxTools.

## Results

### Study approach, quality control, and comparison of metrics

To evaluate the clinical utility of *ex vivo* drug sensitivity profiling in AML, we systematically performed drug screens on bone marrow or peripheral blood samples from patients that were obtained at the time of diagnosis (Fig. 1A and Suppl. Table S1). For high-throughput curve-fitting and multiparametric scoring of drug sensitivities, we used the Breeze pipeline, which performs Hill curve fitting to compute several drug sensitivity metrics, such as the EC50, TEC50, and DSS1, DSS2 and DSS3^22,25^. In addition to these metrics, we calculated a raw AUC metric (rAUC) by computing a step-wise normalized rectangular area under the relative viability dose response along a logarithmic concentration scale (Fig. 1B). Finally, due to potential right skewness in the relative viability scale, we also performed a negative log_2_-transformation of the rAUC, resulting in a weighted-average log_2_ fold-change in cell viability (rAUC-log_2_).

**Figure 1.**
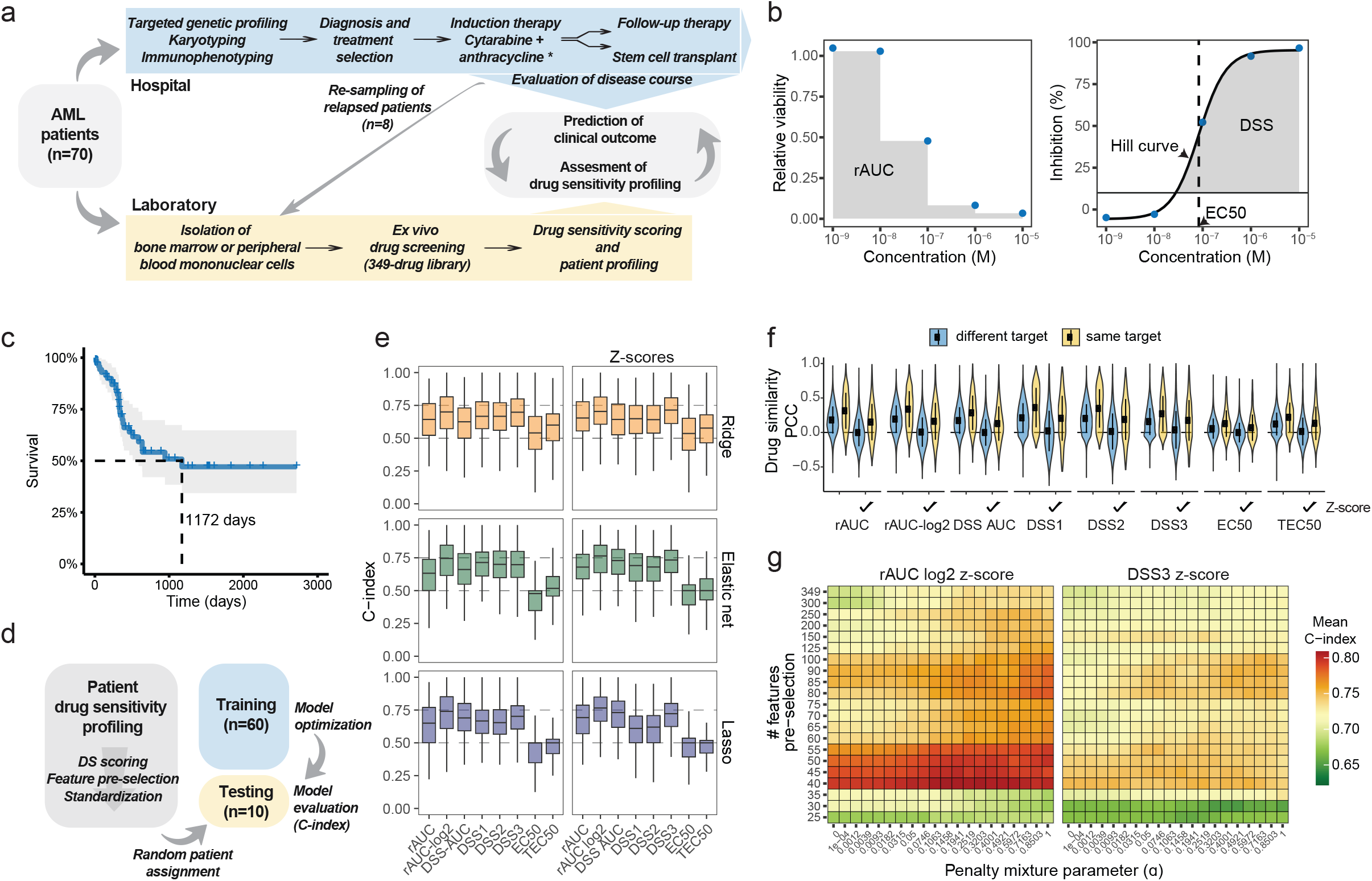
Survival prediction from *ex vivo* drug sensitivity profiles. ***A***, Study overview and workflow. ***B***, Drug sensitivity metric computation. The gray areas indicate how the rAUC and DSS are calculated. ***C***, Survival curve of the study cohort. Ticks indicate censoring. ***D***, Machine learning routine for testing survival prediction from different drug sensitivity metrics and data processing operations. ***E***, C-index results (200 tests) for Cox models trained on different drug sensitivity metrics (*left*) or drug sensitivity z-scores (*right*). ***F***, Pearson correlation coefficients of drug sensitivities between drug pairs having the same or different target. ***G***, Mean test C-index results (50 tests) for Cox models trained on rAUC-log_2_ or DSS3 z-scores with various penalty mixture parameters (□) and feature pre-selection thresholds based on rAUC standard deviations.

To obtain a measure of the quality of the drug screens we computed plate-wise Z’-factors, and analyzed drug sensitivity profile correlations between patients to detect potential outliers (Suppl. Fig. S1). Although some patient-specific plates had a suboptimal Z’-factor (Suppl. Fig. S1A), all plate control averages were at least two standard deviations apart (Suppl. Fig. S1B), and we did not observe any consistent association between plate control noise and patient profile correlations (Suppl. Fig. S1C). However, we did find substantial differences in profile correlations for different metrics, with DSS1-3 and rAUC-log_2_ yielding the best inter-patient correlations (median PCC over 75%), and TEC50 and EC50 giving the worst (Suppl. Fig. S1C-D). Furthermore, intra-patient profile correlations between treatment-naive and relapsed samples were higher than inter-patient correlations (Suppl. Fig. S1D). Comparison of the different metrics revealed that low-confidence Hill curve-fits reported by Breeze tend to be associated with low-sensitivity dose responses (EC50 >10^−7^ μM) (Suppl. Fig. S2A), as well as with responses yielding lower correspondence between AUC metrics that are based either on different regression models (Hill versus LOESS; DSSs) or on raw measurements (rAUCs). Despite this, the overall correlation between the different metrics was high, indicating that inferences made from the drug screen dataset were sufficiently robust for these analysis methods.

### Assessing the predictivity of drug sensitivity metrics

Median survival in the patient cohort was 1172 days (Fig. 1C). To assess *ex vivo* drug sensitivity profiling in predicting patient survival, we compared several regularized Cox models trained on the different drug sensitivity metrics (Fig. 1D). Predictive accuracy was assessed using a C-index over multiple randomized assignments of patient test data (Fig. 1D). We also tested different regularization types, as Ridge and Elastic net regression tend to perform better than Lasso when there are grouped correlations between features (which is expected for drugs with similar modes of action)^34^. AUC-based metrics generally outperformed EC50 and TEC50 in predicting patient survival (Fig. 1E). Interestingly, the DSS-AUC, which had a strong linear correlation with the rAUC (Fig. S2A), improved the survival prediction for Elastic net and Lasso models, suggesting that de-noising dose-response data using a LOESS curve-fit has a beneficial effect (Fig. 1E, left). This was also the case for the DSS scores for all three model types, with DSS3 outperforming DSS1 and DSS2 and resulting in lower overfitting (Fig. 1E and Suppl. Fig. S3A). Strikingly, log-transforming the rAUC resulted in a substantial improvement in prediction accuracy, performing at least as good as DSS3 and better than DSS-AUC (Fig. 1E, left), indicating that distributional skewness in relative viabilities has a negative impact on appropriate drug sensitivity scoring.

To counter potential batch effects, we standardized the drug sensitivity distribution for each patient (Suppl. Fig. S2B). This technique transforms the drug sensitivity metrics to a respective z-score, assuming that the major differences in patient distributions mainly reflect technical influences and that only changes in the relative magnitude of specific drug sensitivities are relevant. With the exception of DSS1 and DSS2, this operation further improved the prediction accuracy for the AUC-based metrics, with rAUC-log_2_ yielding the best performance with a median C-index of 76% for both Elastic net and Lasso, whereas DSS3 and AUC-DSS yielded a median C-index of 73% for Elastic net (Fig. 1E, right). In contrast, patient standardization of DSS1 and DSS2 caused a slight reduction in predictivity that was most profound for models trained with a Lasso penalty (Fig. 1E). To better understand these effects, we measured the inter-drug profile correlations, which revealed library-wide multicollinearity that was removed when standardizing the metrics, but maintained to a certain degree for drug pairs targeting similar proteins (Fig. 1F and Suppl. Fig. S2C-D). Here, DSS1 and DSS2 showed a slightly stronger correlation than DSS3 and other AUC metrics.

Scaling the drug distributions (Suppl. Fig. S2B) to equalize the penalty of drugs with differences in response variance caused a strong reduction in prediction accuracy using rAUCs, while retaining or slightly improving the prediction accuracy using DSS1 and DSS2 (Suppl. Fig. S3B). This suggests that the strict processing of the DSS de-noises the data but also removes informative variability, and that patient-wise standardization of other metrics may improve predictions by removing batch-related multicollinearity and noise.

Given the large number of drugs relative to the number of patients, we also tested the effect of pre-selecting features ranked by their standard deviations as well as a greater range of Elastic net penalties. These operations, in particular feature pre-selection, improved the survival predictions further, with rAUC-log_2_ yielding C-index averages over 80% for several Elastic net models (Fig. 1G and Suppl. Fig. S3D-F). Altogether, these results show that there is high prognostic value in patients’ ex vivo drug sensitivity profiles.

### Removal of confounding factors in *ex vivo* drug sensitivity profiles

Next, we performed principal component analysis (PCA) to identify potential confounding factors that may cause variability in the datasets, such as artefacts associated with curve-fitting, batch covariates, instrument and sample type, and biological/clinical covariates based on genetic and diagnostic phenotypes.

The first principal component identified over 20% of the total variance (Fig. 2A, left), and was strongly associated with curve-fit non-responders (>75%, Fig. 2B, left). Moreover, noise in plate controls and curve-fit error had strong associations with the first two to four components of DSS2 and DSS3, whereas rAUCs were more evenly affected (Fig. 2B, left). Strikingly, for any drug sensitivity metric the first component was almost devoid of any association with biological/clinical covariates, suggesting that the major cause of variability in drug sensitivities stems from patient response averages, noise and potential curve-fit issues.

**Figure 2.**
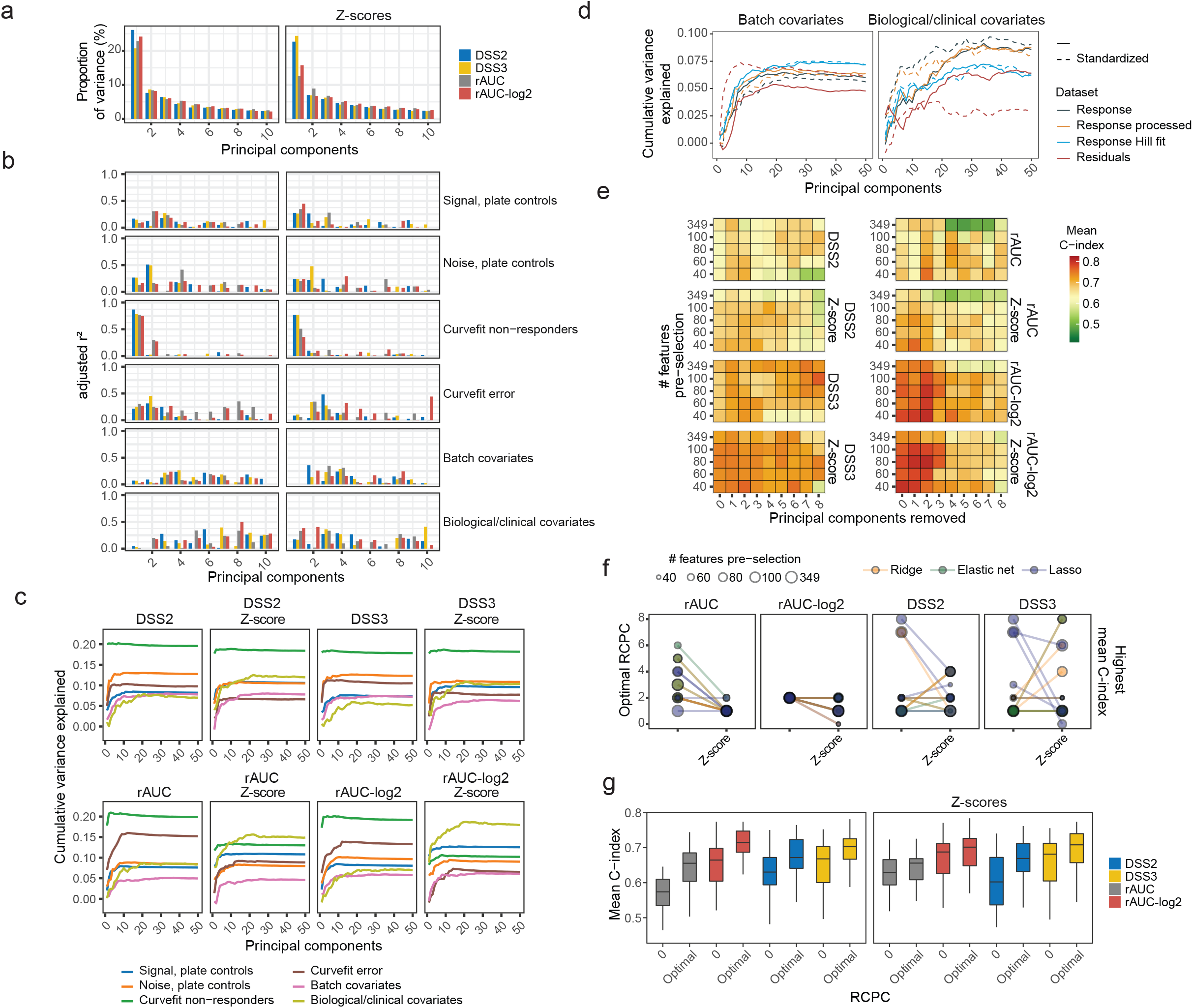
Exploring confounding factors with PCA. ***A***, Percent variance explained by principal components for different drug sensitivity metrics and z-scores. ***B***, Principal component variance explained by different patient sample characteristics, for different drug sensitivity metrics and z-scores. ***C***, Cumulative variance explained by different patient sample characteristics, for different drug sensitivity metrics and z-scores. ***D***, Cumulative variance explained by batch or clinical/biological covariates, for different dose-response datasets. ***E***, Mean test C-index results (50 tests) for Lasso survival models trained on different drug sensitivity metrics or z-scores with various feature pre-selection thresholds (based on rAUC standard deviations), and different numbers of principal components removed. ***F***, Number of components removed to achieve the highest mean test C-index for different drug sensitivity metrics or z-scores shown in Fig. 2E and Suppl. Fig. S4B. ***G***, Mean test C-index results (50 tests) for Lasso survival comparing zero or the optimal number of components removed for 50 datasets generated under weighted random sampling of features, for different drug sensitivity metrics or z-scores (shown in Suppl. Fig. S4E).

These effects were rectified to some extent by standardizing the drug sensitivities per patient, particularly for rAUC and rAUC-log_2_ (Fig. 2A-C). It also increased the variance explained by biological/clinical covariates, while decreasing the effect of other sample characteristics, such as number of patient non-responders (Fig. 2B and C). Surprisingly, DSS2 and DSS3 were most susceptible to batch covariates and noise in plate controls (Fig. 2C), and PCA analysis of the raw, processed, curve-fitted and residual dose-response data from Breeze revealed that the curve-fitting is inherently sensitive to batch variability, introducing biases that confound the information in DSS-based patient profiles (Fig. 2D).

To remove confounding factors that interfere with accurate prediction of patient survival, we performed a Removal of Confounding Principal Components (RCPC) procedure^35^. For all non-standardized datasets, removing at least one or two principal components improved prediction accuracy (Fig. 2E-F and Suppl. Fig. S4), and although all metrics benefitted from RCPC, rAUC-log_2_ improved most in average prediction accuracy (Fig. 2G). Furthermore, removing a greater number of principal components resulted in a sharp decline in prediction performance of rAUCs (Fig. 2E), indicating loss of valuable information about patient survival. Finally, standardization of the rAUCs resulted in a significant decrease in component subtractions needed to achieve optimal performance (Fig. 2F), and a lower improvement in prediction accuracy from RCPC (Fig. 2G). These results indicate that processed *ex vivo* drug screening data can contain consequential confounders that may bias the dose-response curve fit and drug sensitivity scoring, and that standardization or RCPC can be used as a straightforward and reliable method for de-confounding the data.

### Integration with AML biomarkers and treatment response predictions

We next compared the effectiveness of *ex vivo* drug profiling in predicting treatment outcome to the predictiveness of other AML patient data, such as genetic features and/or prognostic clinical features that included ELN risk stratifications, age, sex and WHO’s patient performance status. As expected, due to coverage sparsity the Ridge models performed better on the clinical feature sets (Fig. 3A). The genetic feature set provided the highest prediction with a median C-index of 63%, which is similar to what has been reported previously using much larger cohorts^36^. Combining the different clinical feature sets did not improve predictivity, most likely due to redundancy in information (Suppl. Fig. S5A). In contrast, combining the clinical prognostic features with either the full standardized rAUC-log_2_ or DSS3 datasets resulted in a median C-index improvement of 3-5% (Fig. 3A). Importantly, only minor effects were observed when combining genetic features with the entire drug dataset. However, we did notice that combining genetic features with a reduced drug dataset using feature pre-selection resulted in an improved combination effect (Fig. 3A, lower panel and Suppl. Fig. S5B), indicating that reducing the drug library may improve complementarity with the genetic data by removing redundant information.

**Figure 3.**
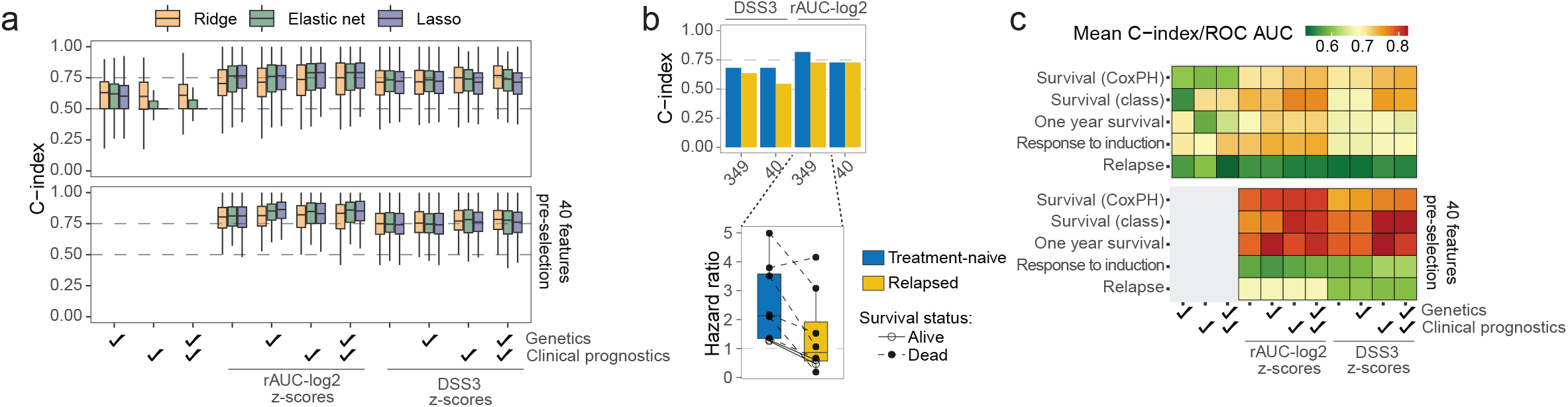
Integration with AML biomarkers for clinical outcome predictions. ***A***, Test C-index results (200 tests) for Cox models trained on different dataset compositions based on clinical feature sets and rAUC-log2 or DSS3 z-scores. The lower panel represents pre-selection of 40 features based on rAUC standard deviations. ***B***, Test C-index and predicted hazard ratio (relative to population-median) on samples from treatment-naive and relapsed patients, for Lasso survival models trained on rAUC-log2 or DSS3 z-scores with or without feature pre-selection. ***C***, Comparing Ridge model mean test C-index from (*A*) with mean test ROC-AUC scores for classification of various binarized clinical outcomes. Survival (class) represents the classification of overall survival irrespective of survival times. Dataset compositions are identical to (*A*).

To evaluate the prognostic versatility and robustness of *ex vivo* drug profiling, we also tested the prediction on relapsed samples and forecasting potential of other clinical outcomes. Models for overall survival trained on treatment-naive samples consistently showed a slight reduction in prediction accuracy on relapsed samples with a reduction in predicted patient hazard (Fig. 3B and Suppl. Fig. S5C-D). This suggests that the cancer dependencies relevant for prognostics of high-risk patients evolve during the course of therapy, which prompted us to test whether *ex vivo* drug screening can predict the response to induction therapy, risk of relapse, and one-year survival. Indeed, *ex vivo* drug sensitivities were capable of predicting induction response and relapse with accuracies similar to that of genetic and clinical prognostic feature sets (Fig. 3C), as well as showing substantial complementary utility. We conclude that *ex vivo* drug profiling can predict the course of disease and treatment outcome.

### Drug sensitivity risk associations

A statistical analysis of the library-wide survival associations revealed that models trained on rAUC-log_2_ yielded more robust model coefficients compared to DSS3 (Fig. 4A and Suppl. Fig. S6A). Nonetheless, different models showed overall good correlation in their learned coefficients (Fig. 4B and Suppl. Fig. S6B-C), where the strongest survival associations also corresponded to high contributions to a model’s prediction accuracy (Fig. 4C and Suppl. Fig. S6D). Given that many compounds in the library have overlapping targets, there was substantial redundancy in single-drug withdrawals with respect to model performance (Fig. 4C). In-depth investigation of specific drug responses that were significantly associated with survival revealed that *ex vivo* sensitivity to anthracyclines predicted a favorable outcome (Fig. 4A). Daunorubicin, one of the major components of 7+3 chemotherapy, had one of the most significant associations with survival, while Idarubucin also had a favorable (albeit statistically non-significant) survival association. Epirubicin, another anthracycline used in treatment of several types of cancers, and the anthracycline analog Mitoxantrone, were also significantly associated with survival (Fig. 4A). Interestingly, we found that *ex vivo* sensitivity to the Bcl-2/Bcl-XL/Bcl-w inhibitor ABT-263 (Navitoclax) was strongly associated with increased risk (Fig. 4A). ABT-263 is an experimental drug with a mechanism of action similar to the more selective Bcl-2 inhibitor Venetoclax, which has been reported to significantly improve survival of AML patients who are ineligible for standard chemotherapy (please note that at the time of sample collection, Venetoclax had not yet been approved for AML therapy, which is why it was absent from the drug library).

**Figure 4.**
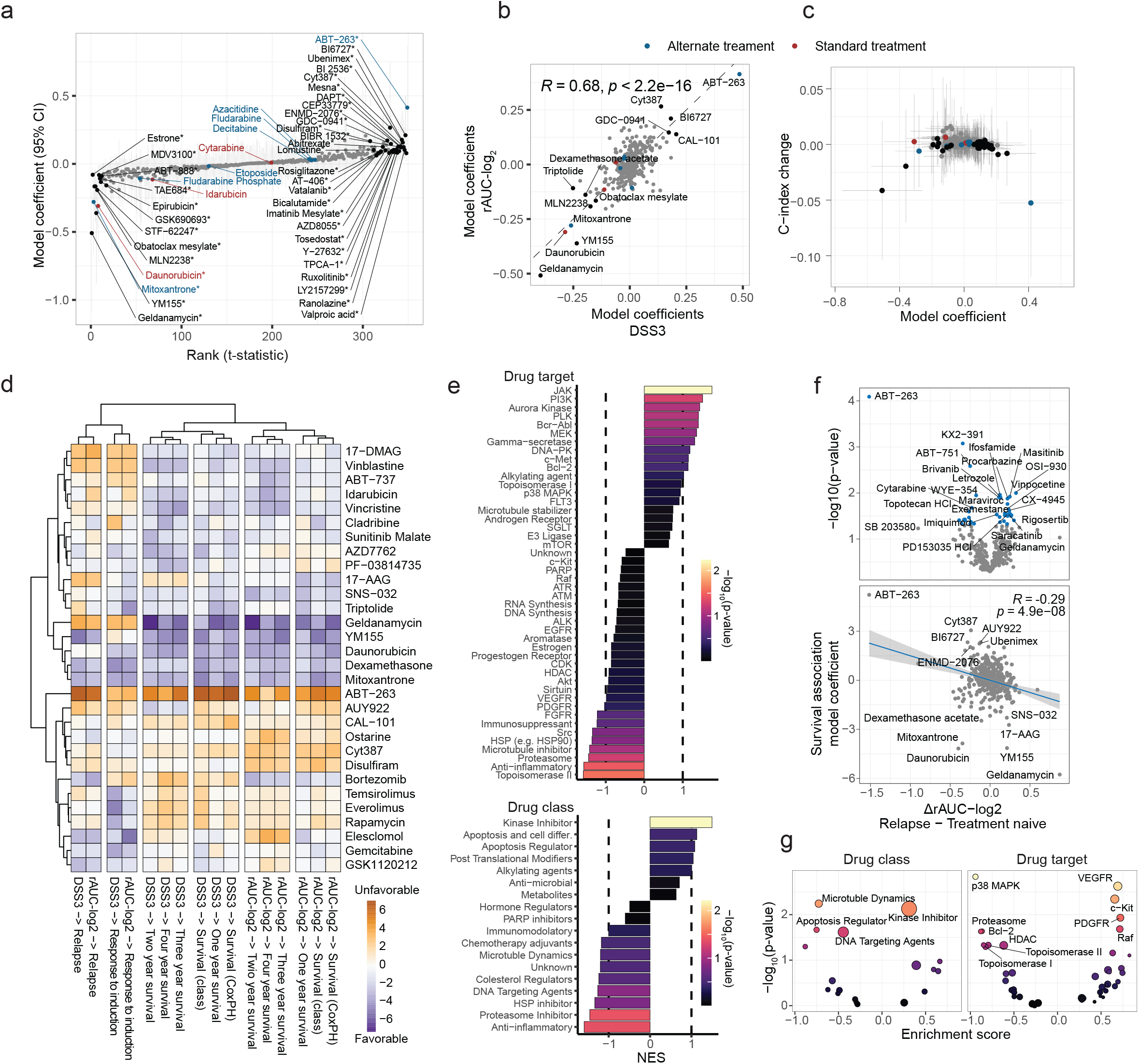
Clinical associations of *ex vivo* drug sensitivities. ***A***, Bootstrapped Ridge survival coefficients representing risk association of AUC-log_2_ z-scores for 349 drugs. Gray vertical bars indicate 95% confidence intervals for each individual compound. Significance (*) was determined when 95% of the bootstrapped coefficients did not include or cross zero. ***B***, Correlation between estimated Ridge survival coefficients for rAUC-log_2_ z-scores or DSS3 z-scores. The drugs with the strongest coefficients in both models are labeled. ***C***, Association between mean survival coefficients and mean test C-index change (50 tests) in response to drug withdrawal using Ridge regression on rAUC-log_2_ z-scores. Horizontal and vertical bars indicate standard deviations. Standard or alternative AML treatment drugs are color-coded in (*A-C*). ***D***, Clustering of standardized Ridge coefficients from models against different clinical outcomes. ***E***, Drug target or drug class risk associations based on directional drug-set enrichment on ranked Ridge survival coefficients (for rAUC-log_2_ z-scores). ***F***, Differential drug sensitivity (for rAUC-log_2_ z-scores) between samples from relapsed and treatment-naive patients, with paired t-test p-values (*upper*) and association with Ridge survival coefficients (*lower*). Blue dots indicate significance. ***G***, Drug target or drug class association with differential sensitivity, using directional drug-set enrichment on differential drug sensitivities from (*F*).

Hierarchical clustering of learned risk associations against different clinical outcomes revealed interesting correlations (Fig. 4D). For instance, sensitivity to the mTOR inhibitors Rapamycin, Everolimus, and Temsirolimus was associated with initial positive response to induction therapy, but also with increased risk of relapse and decreased long-term survival. This implies that dependence on mTOR may be an early prognostic marker for acquired resistance to chemotherapy. Furthermore, enrichment analysis of drug targets associated with Cox model coefficients demonstrated a library-wide risk association of sensitivity to kinase inhibitors, including JAK and PI3K signaling (Fig. 4E). Analysis of the differential sensitivity between treatment-naive and relapsed samples indicated a further increase in kinase inhibitor sensitivity, including multiple receptor tyrosine kinases (Fig. 4F-G). Conversely, sensitivity to topoisomerase II-targeting chemotherapeutics in treatment-naive samples, which was associated with a favorable outcome, was decreased in samples from relapsed patients (Fig. 4E-G). Consistent with previous studies^37,38^, *ex vivo* sensitivity to agents that target anti-apoptotic proteins, particularly ABT-263, was strongly reduced in relapsed samples (Fig. 4F), indicating an increased apoptotic threshold or development of cross-resistance. This was also accompanied with a library-wide negative correlation trend between differential relapse sensitivity and risk-association in treatment-naive samples, where several prognostic drug markers for adverse response to standard treatment diminished in sensitivity after relapse (Fig. 4F, lower). We conclude that *ex vivo* drug sensitivity profiles can serve as prognostic markers and may provide novel insight into relevant cancer dependencies and mechanisms of disease progression.

### Drug sensitivity-based risk stratification

Finally, to investigate the relationship between different drug responses and their contribution to survival forecasting, we used the drug sensitivity profiles to cluster the patients into different risk groups. In contrast with previous studies comparing drug sensitivity metrics, we could only detect minor differences in clustering fidelity based on a Rand index measuring the agreement between curated drug assignment and hierarchical clustering (Suppl. Fig. S7A-B)^25^. However, congruent with the previous findings in this study, clustering of patients improved using DSS or rAUC-log_2_ compared to rAUC, as well as with patient standardization (Suppl. Fig. S7C). To assess drug profiles with the highest relevance for risk assignment, we weighted the rAUC-log_2_ drug sensitivity data with learned Lasso model coefficients, thus shrinking the drug dimension and transforming the scores to relative risk contributions (Fig. 5A). Although this reduced the agreement with clusters of genetic markers slightly, it greatly improved patient risk group stratification (Fig. 5B and Suppl. Fig. S7C-D). Stratifying the patient cohort into six groups, clear patterns in relative drug sensitivity profiles emerged that aligned with different survival responses (Fig. 5). For instance, intermediate sensitivity to Daunorubicin combined with high sensitivity to ABT-263 represented a large sub-group with a high relapse rate and very short-term survival. Conversely, increased Daunorubicin sensitivity for high-responders to ABT-263 was associated with intermediate risk, but when accompanied by low sensitivity to ABT-263 it was associated with good prognosis. Furthermore, high sensitivity to Daunorubicin as well as the HSP-90 inhibitor Geldanamycin was associated with positive outcome. Taken together, *ex vivo* drug screening can stratify patients in risk groups.

**Figure 5.**
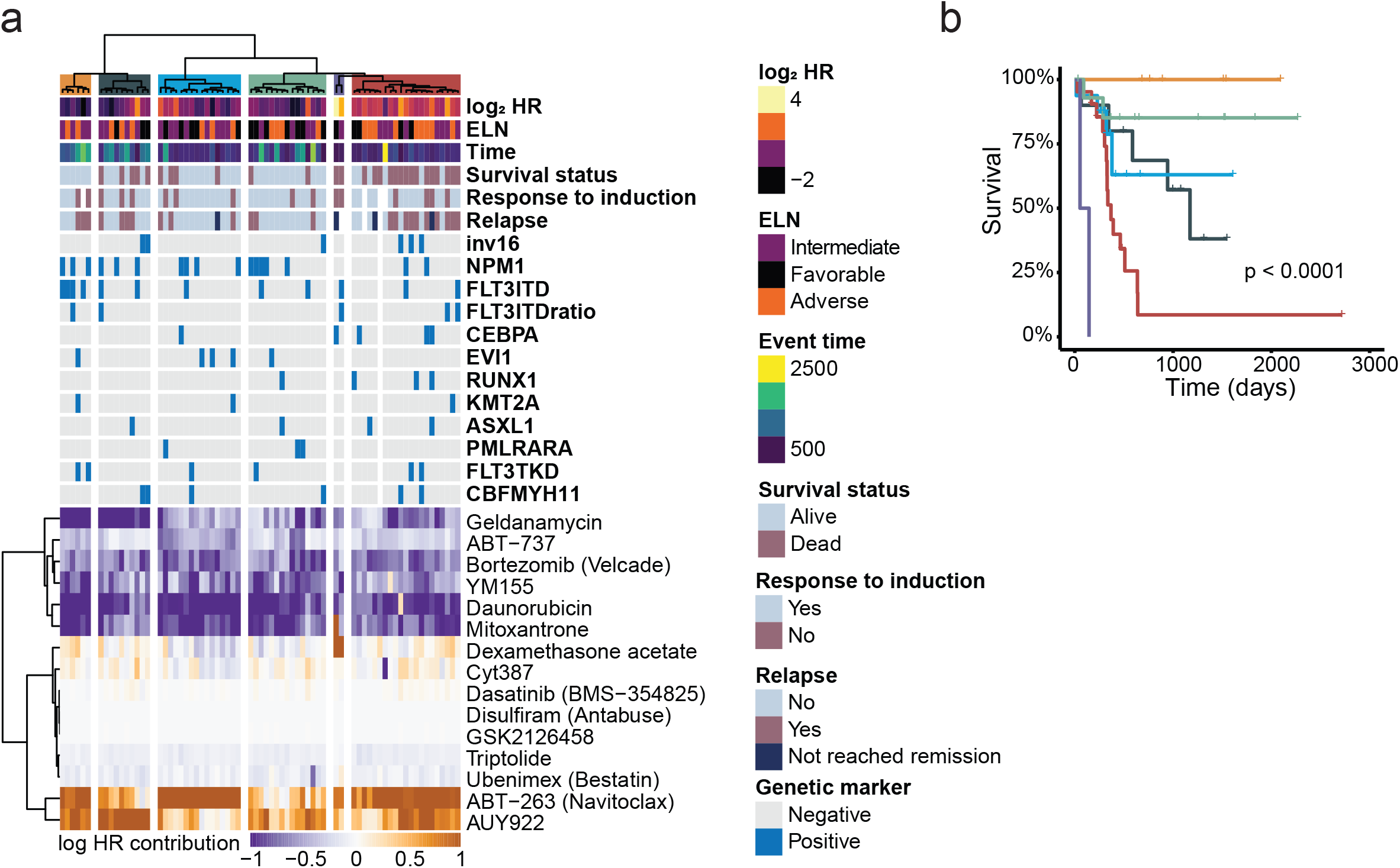
Patient risk group stratification based on *ex vivo* drug sensitivities. ***A***, Hierarchical clustering of survival-associated drug sensitivities. Patients are clustered along the columns and drugs are clustered along the rows. rAUC-log_2_ z-scores were multiplied with Lasso survival coefficients for each drug to generate log-HR contribution per individual drug sensitivity, and zero-sum rows were removed. Patients are color annotated for mutations, sex, response to induction, relapse, survival status, event time, prognosis, and predicted log-HR (relative to population-median). ***B***, Stratification of patients into six risk groups with their respective survival curves, corresponding to the six color-coded columns indicated at the top of the clusters in panel *A*.

## Discussion

In this retrospective study, we used *ex vivo* drug sensitivity screening to predict the survival of AML patients subjected to standard chemotherapy. An important topic that has been the subject of much discussion is how to appropriately score drug sensitivities in large-scale drug screens. Although sigmoid-constrained curve fitting and response scoring is a valuable tool in large-scale drug screens, it is inherently sensitive to dose coverage in the dynamic range, and model parameters tend to be unreliable^23,28,39^. A common approach is to only consider the area under inhibitory response curves that cross a certain threshold, exemplified by the DSS^25^. We observed that this compression of drug response variability resulted in zero-inflated data with substantial loss of information. Furthermore, covariances in curve-fitted inhibition data were more strongly affected by batch effects than covariances in unprocessed dose-response data. We showed that log-transformation of rAUCs outperformed other methods in scoring drug sensitivities, suggesting this is a useful strategy to counter skewness and noisy measurements in large-scale *ex vivo* drug sensitivity screens.

In our study, we used a drug library that not only includes standard chemotherapeutics but also a wide variety of other compounds, and identified informative ‘drug sensitivity fingerprints’ that may not only help guide treatment choice but that can also provide valuable prognostic information. Importantly, merging the data from drug sensitivity profiling with information from genetic biomarkers did not further improve the predictivity of drug profiling alone unless a large number of drugs were removed. This suggests that predictive information in genetic profiles is already captured by *ex vivo* drug sensitivities, and indicates that *ex vivo* drug profiling can be further developed into a prognostic tool through rational design of focused drug libraries that fully cover cancer dependencies. Examples of drugs that should be included in such next-generation libraries for risk stratification are BCL2, mTOR and HSP90 inhibitors, which, as already discussed above, are clearly associated with risk. Furthermore, we found that the five top-ranking drugs associated with high risk include three JAK2 inhibitors and one STAT3 inhibitor, which is consistent with previous findings that high JAK2/STAT3 activity causes resistance to chemotherapy^40–42^. We are currently analyzing data from other large-scale *ex vivo* drug screening efforts, such as the FIMM and BEAT-AML datasets^43–45^, which we expect will identify additional compounds associated with risk.

Although progress has been made to update recommendations for AML risk stratification and treatment guidelines, prediction accuracies in large and heterogeneous cohorts remain modest^4,36^. One reason for this is that AML is in part driven by non-genetic alterations^46,47^, including epigenetic alterations and rewiring of metabolic pathways and signaling circuits, which are not easily detected by more conventional methods, such as genomics and transcriptomics. Functional *ex vivo* drug profiling may be better suited for identifying such cancer dependencies^44,45^. Thus, with further research we envision that *ex vivo* drug profiling can be a useful tool in the clinical decision-making process to stratify patients into treatment groups.

We conclude that *ex vivo* drug profiling reveals cellular dependencies leading to chemoresistance and cancer progression and robustly predicts patient survival and response to induction therapy.

## Acknowledgments

We would like to thank all patients who participated in this study, as well as patient representatives from the Norwegian Blood Cancer Society (Blodkreftforingen), members of the Enserink and Knævelsrud laboratories for fruitful discussions, and the PERCATHE and PINpOINT environments for feedback on the study. We also thank Dr. M. Zucknick for suggesting the drug withdrawal test. This study was supported by grants from the Norwegian Health Authority South-East, grant numbers 2017064, 2018012 and 2019096; the Norwegian Cancer Society, grant numbers 182524 and 208012; and the Research Council of Norway through its Centers of Excellence funding scheme (262652) and through grants and 261936, 294916 and 314811.

## Author contributions

Conceptualization: ANA and JME. Project supervision: ANA and JME. Data analysis: AMB and ANA. Prepared figures and tables: AMB and ANA. Performed drug screens: PAD with assistance from LP, DST, LB, and JF. Diagnosed, sampled and treated patients: YF and AL. Wrote the manuscript: AMB, ANA, JME, YF, AL. Extracted patient data from records: MS, PAD, AMB.

## Data Availability Statement

All data are freely available upon request.

## Supplemental Figure Legends

**Figure S1. Drug screen quality controls. *A***, Z’-factor for DMSO and BzCl plate controls per patient. ***B***, Differential z-score for DMSO and BzCl plate controls per patient. ***C***, Mean intra-patient drug sensitivity profile correlation per patient for different drug sensitivity metrics. ***D***, Correlations in drug sensitivity profiles, either between patients or for the same patient before and after relapse.

**Figure S2. Drug sensitivity metric similarities. *A***, Correlations between the different drug sensitivity metrics. High and low confidence Hill curve fits reported by Breeze are color-coded. ***B***, Standardization procedures used in the study. Drug sensitivity z-scores were generated by standardizing each patient distribution. Drug sensitivity scaling was done by standardizing each drug distribution. ***C, D***, Correlations between drugs with a similar mechanism of action, with drug sensitivity metrics in (*C*), and drug sensitivity z-scores in (*D*).

**Figure S3. Survival prediction from *ex vivo* drug sensitivity profiles. *A***, Training C-index results (200 tests) for Cox models trained on different drug sensitivity metrics (*left*) or drug sensitivity z-scores (*right*). ***B***, Test C-index results (200 tests) for Cox models trained on different drug sensitivity metrics (*left*) or drug sensitivity z-scores (*right*) with feature scaling (drug-wise standardization). ***C***, Overview of feature pre-selection procedure and penalty type testing. ***D***, Mean test C-index results (50 tests) for Cox models trained on rAUC, rAUC-log_2_, or DSS3 z-scores with various penalty mixture parameters (□) and feature pre-selection thresholds based on standard deviations of the respective drug sensitivity metric (rAUC, rAUC-log_2_, or DSS3). ***E***, Median test C-index results (50 tests) for Cox models trained on rAUC-log_2_ or DSS3 z-scores with various penalty mixture parameters (□) and feature pre-selection thresholds based on rAUC standard deviations. ***F***, Median test C-index results (50 tests) for Cox models trained on rAUC, rAUC-log_2_, or DSS3 z-scores with various penalty mixture parameters (□) and feature pre-selection thresholds based on standard deviations of the respective drug sensitivity metric (rAUC, rAUC-log_2_, or DSS3).

**Figure S4. Supplement to exploration of confounding factors with PCA. *A***, Overview of PCA procedure based on SVD, and reconstruction using RCPC. ***B***, Mean test C-index results (50 tests) for Ridge and Elastic net survival models trained on different drug sensitivity metrics or z-scores with various feature pre-selection thresholds (based on rAUC standard deviations), and different numbers of principal components removed. ***C***, Number of components removed to achieve the highest median test C-index for different drug sensitivity metrics or z-scores in ***2E*** and ***S4B. D***, Highest mean (upper) and median (lower) test C-index results (50 tests) achieved using RCPC on different drug sensitivity metrics or z-scores with various feature pre-selection thresholds and different Cox models. ***E***, Mean test C-index results (50 tests) for Lasso survival models trained on 50 datasets generated under weighted random sampling of features, using different drug sensitivity metrics or z-scores, and different numbers of principal components removed. ***F***, Number of components removed to achieve the highest mean test C-index for different drug sensitivity metrics or z-scores in ***S4E. G***, Number of components removed to achieve the highest median test C-index for different drug sensitivity metrics or z-scores in ***S4E. H***, Median test C-index results (50 tests) for Lasso survival comparing zero or the optimal number of components removed for 50 datasets generated under weighted random sampling of features, for different drug sensitivity metrics or z-scores (in ***S4E***).

**Figure S5. Supplement to integration with AML biomarkers for clinical outcome predictions. *A***, Training C-index results (200 tests) for Cox models trained on different dataset compositions based on clinical feature sets and rAUC-log2 z-scores or DSS3 z-scores. The lower panel represents pre-selection of 40 features based on rAUC standard deviations. ***B***, Test C-index results (200 tests) for Cox models trained on different dataset compositions based on clinical feature sets and rAUC-log2 or DSS3 z-scores with different feature pre-selection cutoffs. ***C***, Test C-index on treatment-naive and relapsed samples, for Ridge or Elastic net survival models trained on rAUC-log2 or DSS3 z-scores with or without feature pre-selection. ***D***, Predicted hazard ratio (relative to population-median) on treatment-naive and relapsed samples, for Ridge, Lasso, or Elastic Net survival models trained on rAUC-log2 or DSS3 z-scores with or without feature pre-selection. Open circles indicate alive, closed circles indicate deceased patients. ***E***, Testing and training ROC-AUC scores for classification of various binarized clinical outcomes using Ridge models trained on different dataset compositions. ‘Survival (class)’ represents the classification of overall survival irrespective of survival times.

**Figure S6. Supplement to clinical associations of *ex vivo* drug sensitivities. *A***,

Bootstrapped Ridge survival coefficients representing risk association of DSS3 z-scores for 349 drugs. The vertical bars indicate the 95% confidence interval. Significance (*) was determined when 95% of the bootstrapped coefficients did not include or cross zero. ***B***, Correlation between estimated Lasso and Elastic net survival coefficients for rAUC-log_2_ z-scores or DSS3 z-scores.The drugs with the strongest coefficients in both models are labeled. ***C***, Association between mean survival coefficients and mean test C-index change (50 tests) in response to drug withdrawal using Ridge regression on DSS3 z-scores. The horizontal and vertical bars indicate the standard deviations, respectively. Standard or alternate AML treatment drugs are color-coded in ***A-C. D***, Drug target or drug class risk associations based on directional drug-set enrichment on ranked Ridge survival coefficients (for DSS3 z-scores).

**Figure S7. Clustering of *ex vivo* drug sensitivities. *A***, Hierarchical clustering of rAUC-log_2_ z-scores. Patients are clustered along the columns and drugs are clustered along the rows. Patients are color annotated as in ***Fig. 5*** for mutations, sex, response to induction, relapse, survival status, event time, prognosis, and predicted log-HR (relative to population-median). ***B***, Agreement between drug class or drug target assignment and hierarchical clustering of drugs with different drug sensitivity metrics and z-scores. The dot shape indicates whether scaling of the drug distributions was performed before clustering. ***C***, Agreement between different patient characteristics and hierarchical clustering of patients with different drug sensitivity metrics and z-scores. The dot color indicates whether the rows were weighted by estimated Lasso survival coefficients for the respective datasets prior to clustering. ***D***, Survival curves for patient groups based on the dendrogram in ***A. E***, Genetic clustering of patients for ***C*** (*lower left*).

## Notes

### Competing Interest Statement

The authors have declared no competing interest.

### Summary of Updates

Manuscript has been rewritten and figures have been replaced with new figures

