## Supplementary figures and images for "Clinical Forecasting using *Ex Vivo* Drug Sensitivity Profiling of Acute Myeloid Leukemia"

### Suppl. Fig. S1

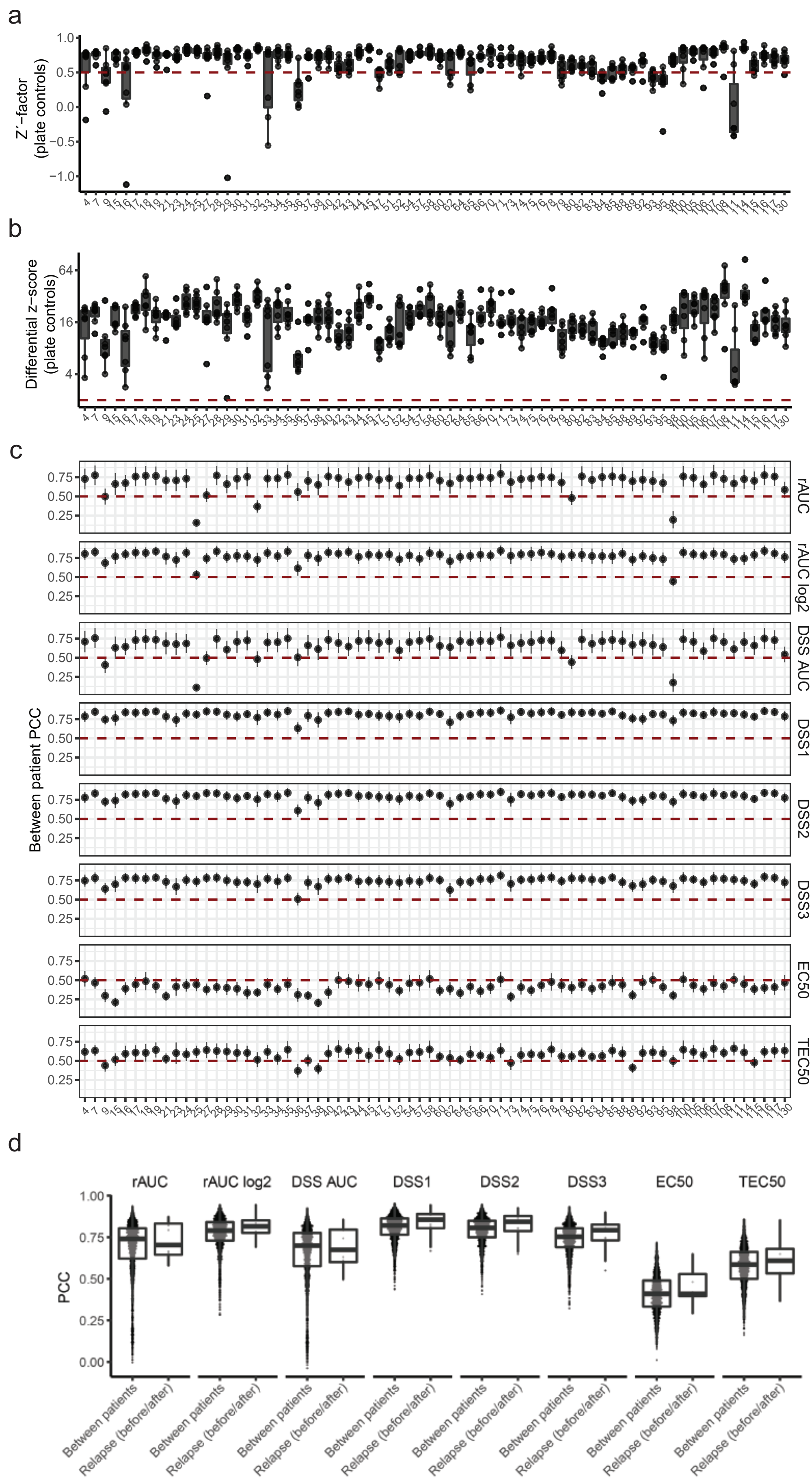

### Suppl. Fig. S2

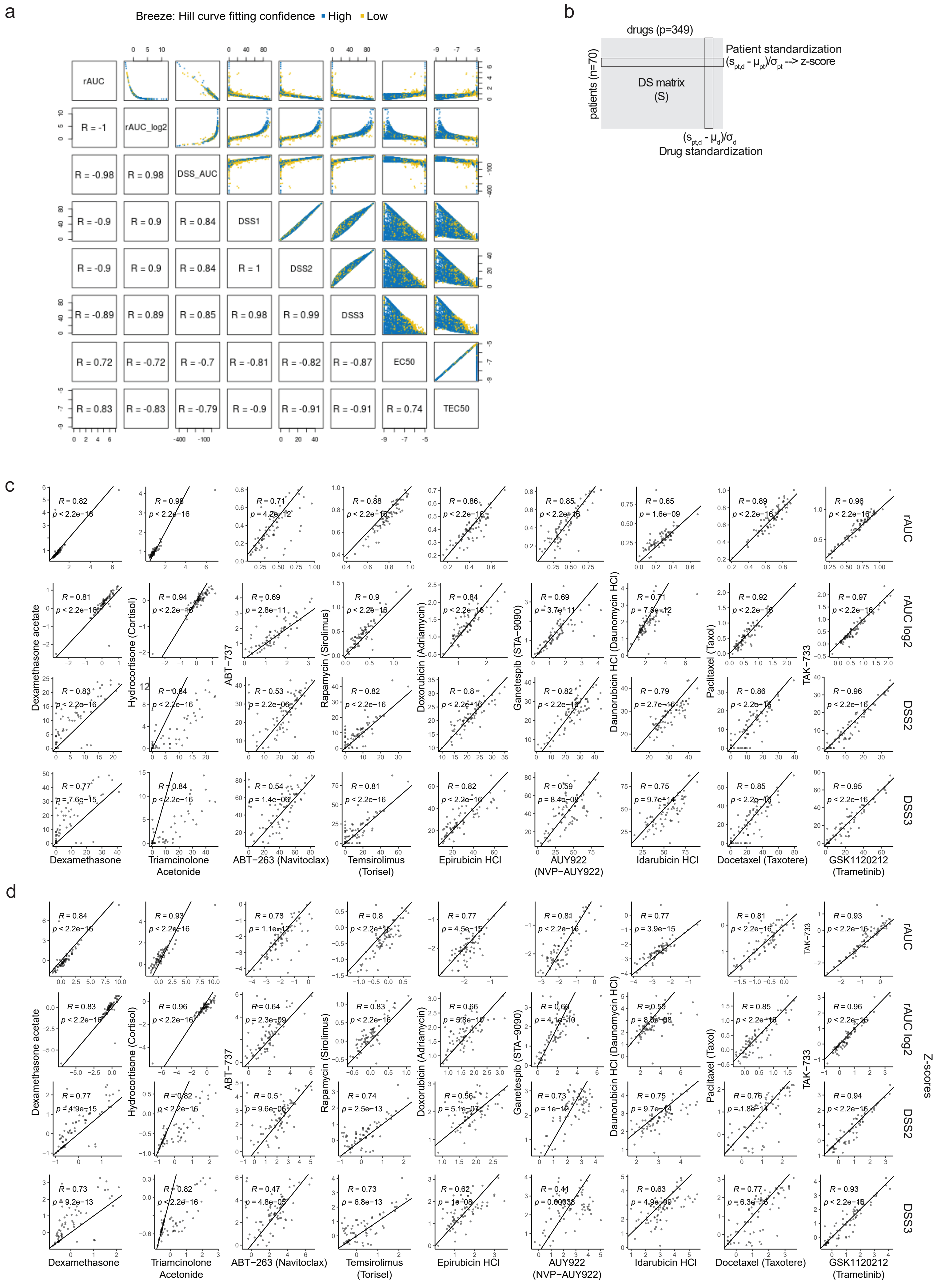

### Suppl. Fig. S3

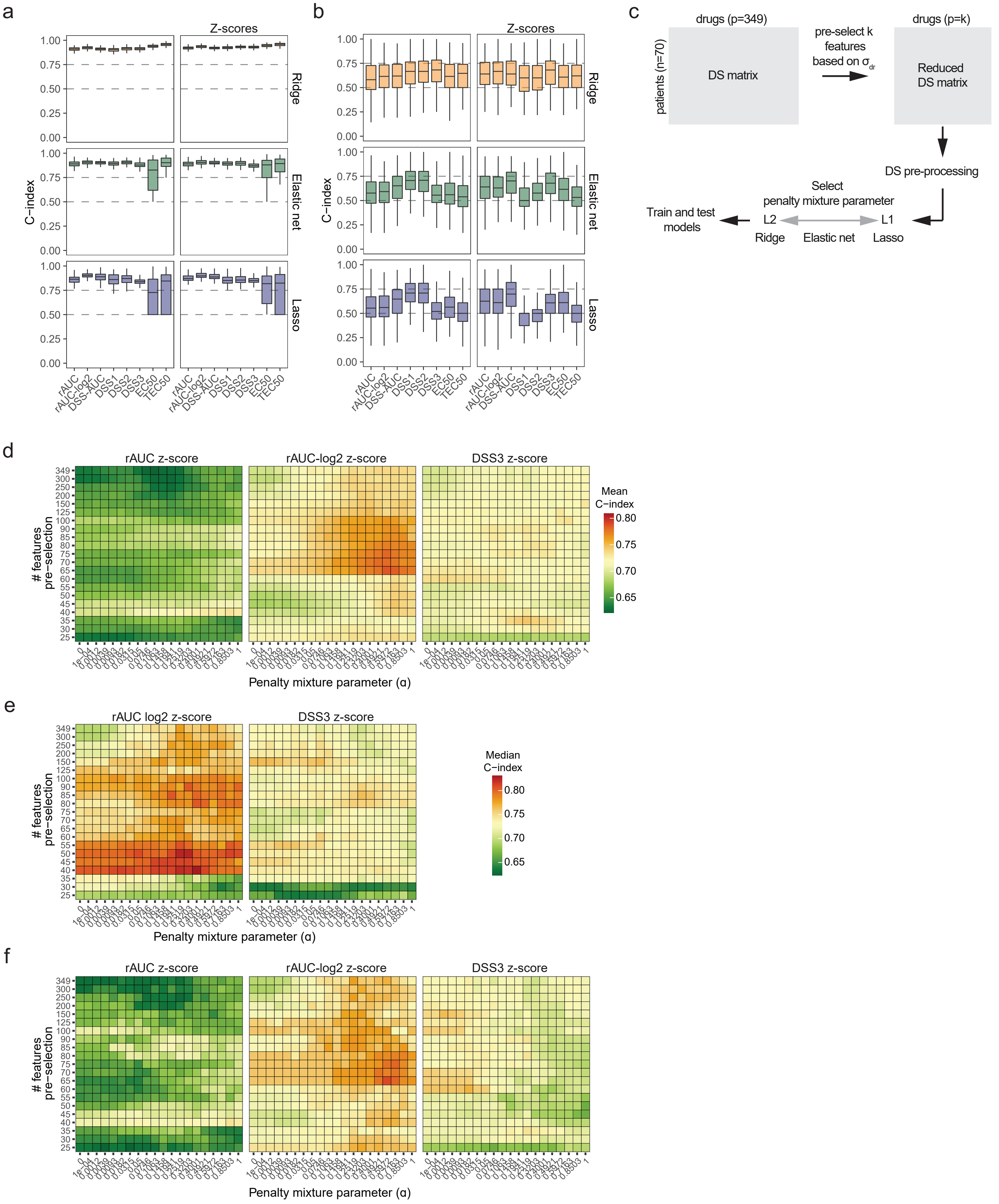

### Suppl. Fig. S4

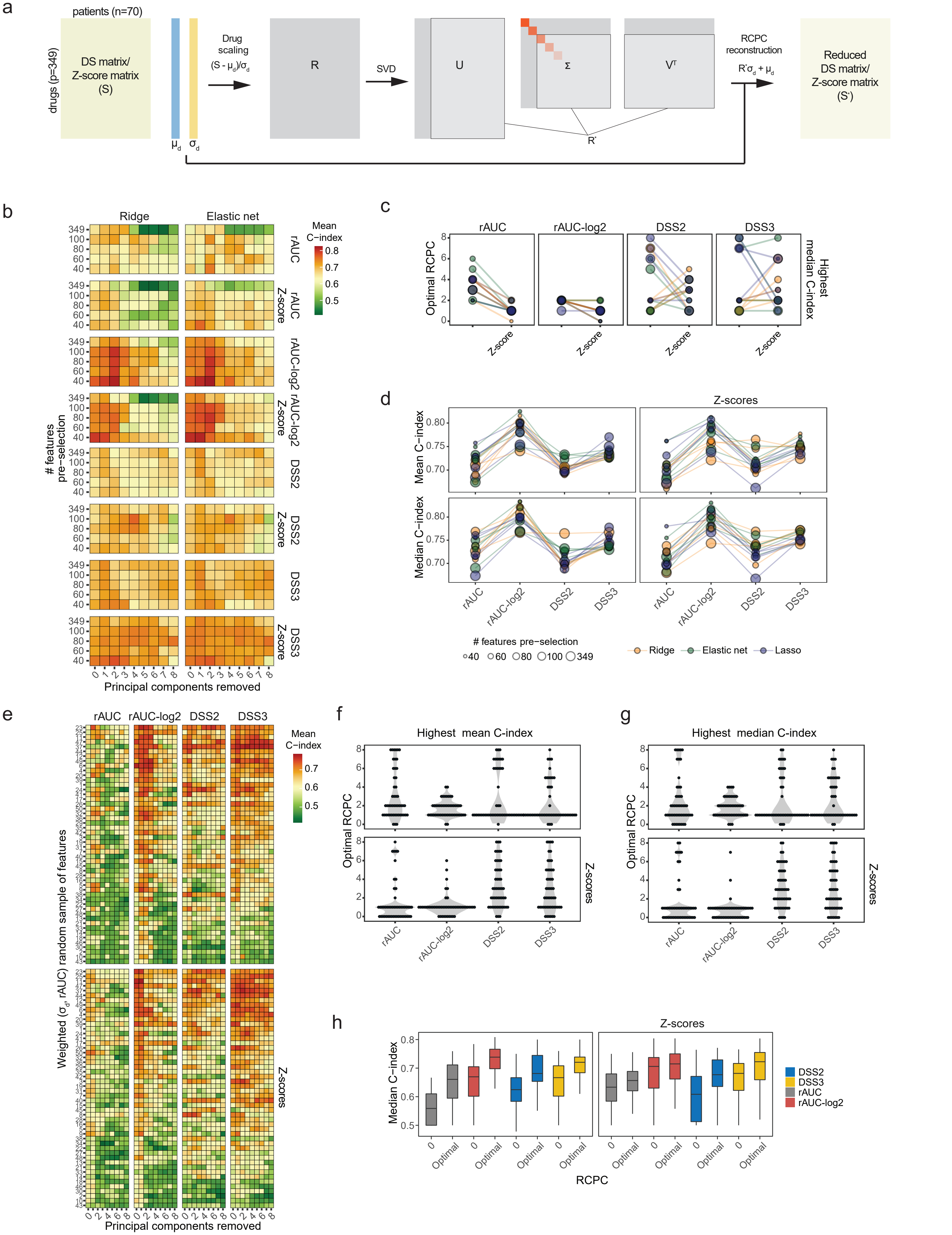

### Suppl. Fig. S5

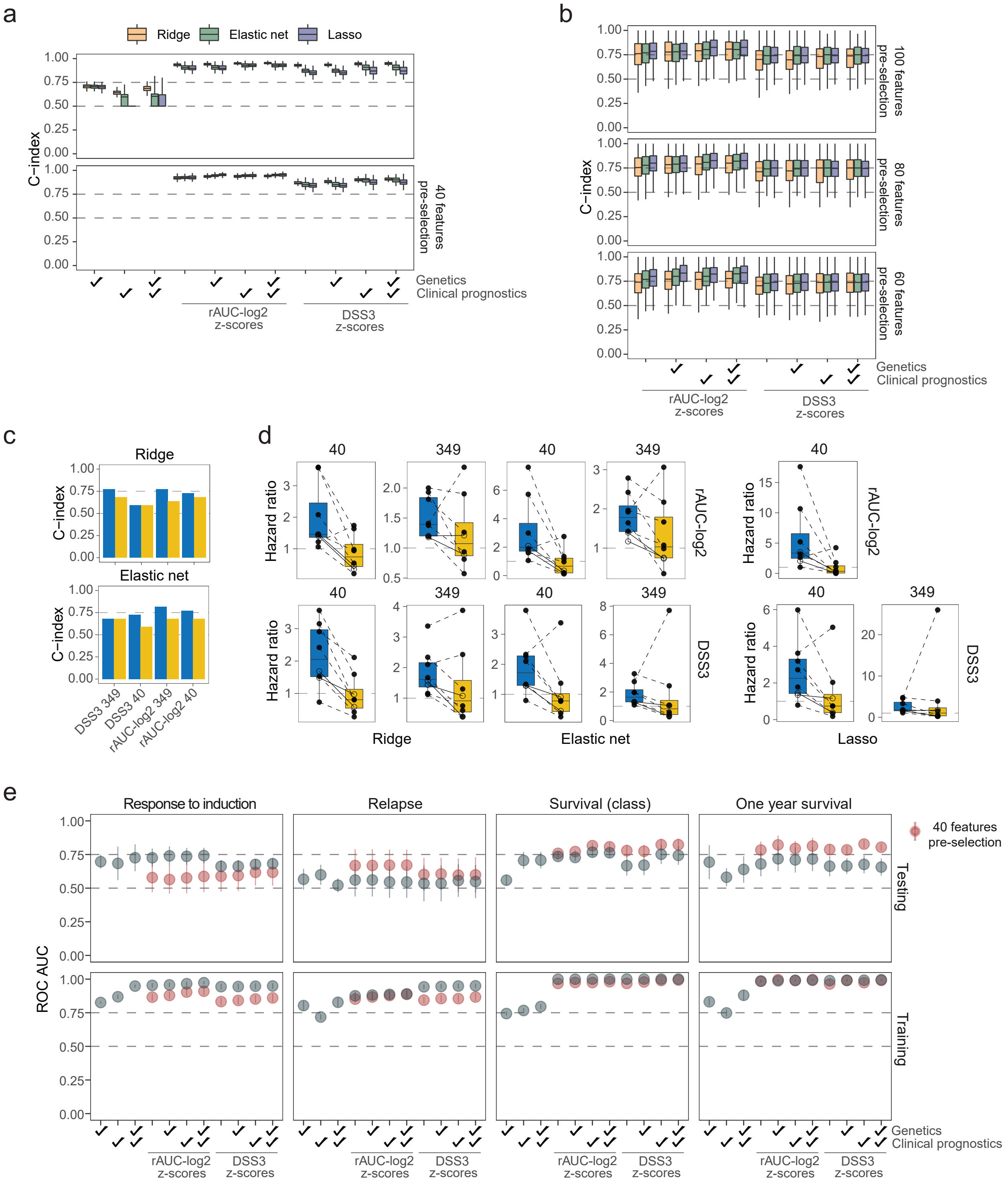

### Suppl. Fig. S6

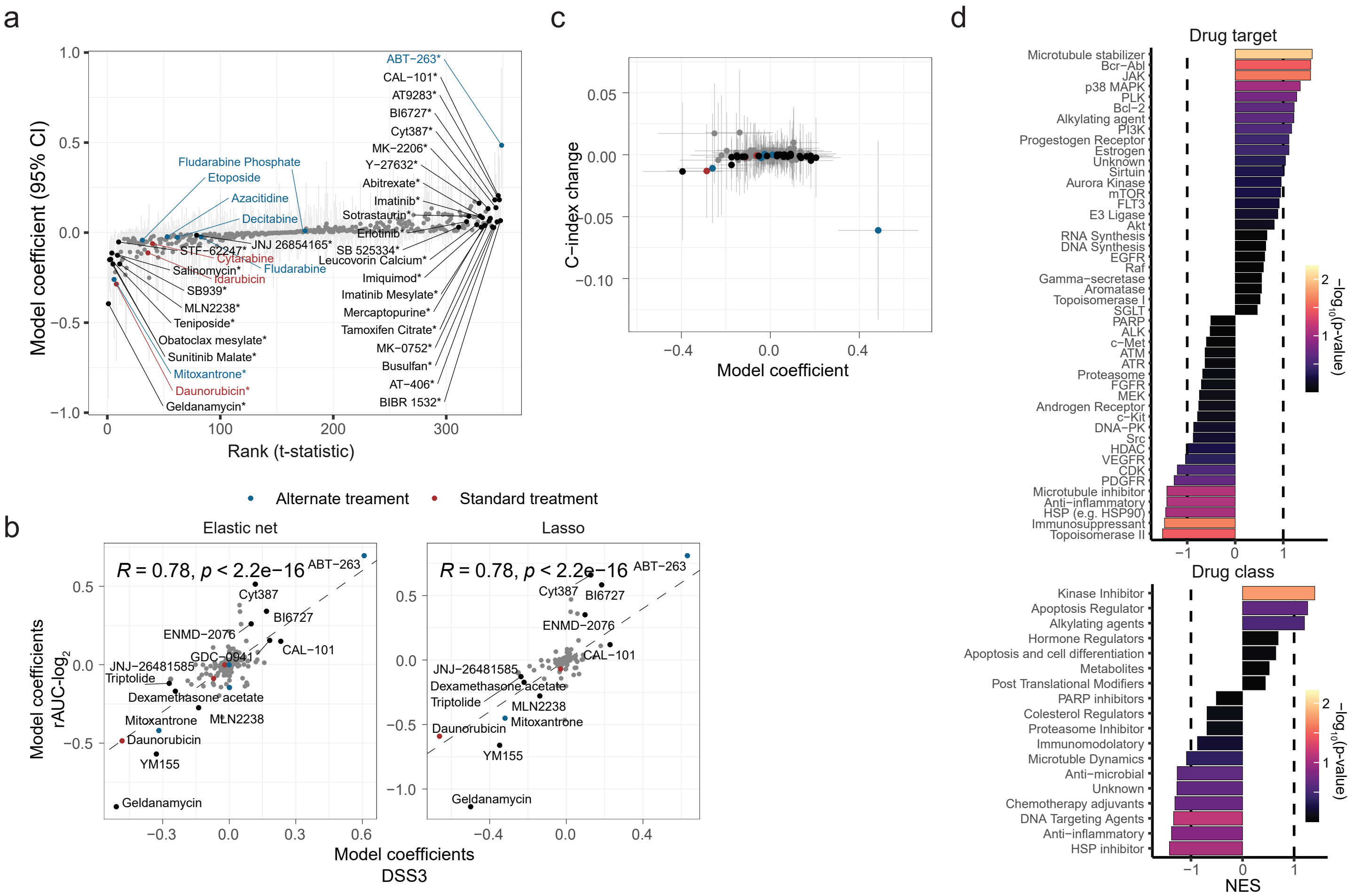

### Suppl. Fig. S7

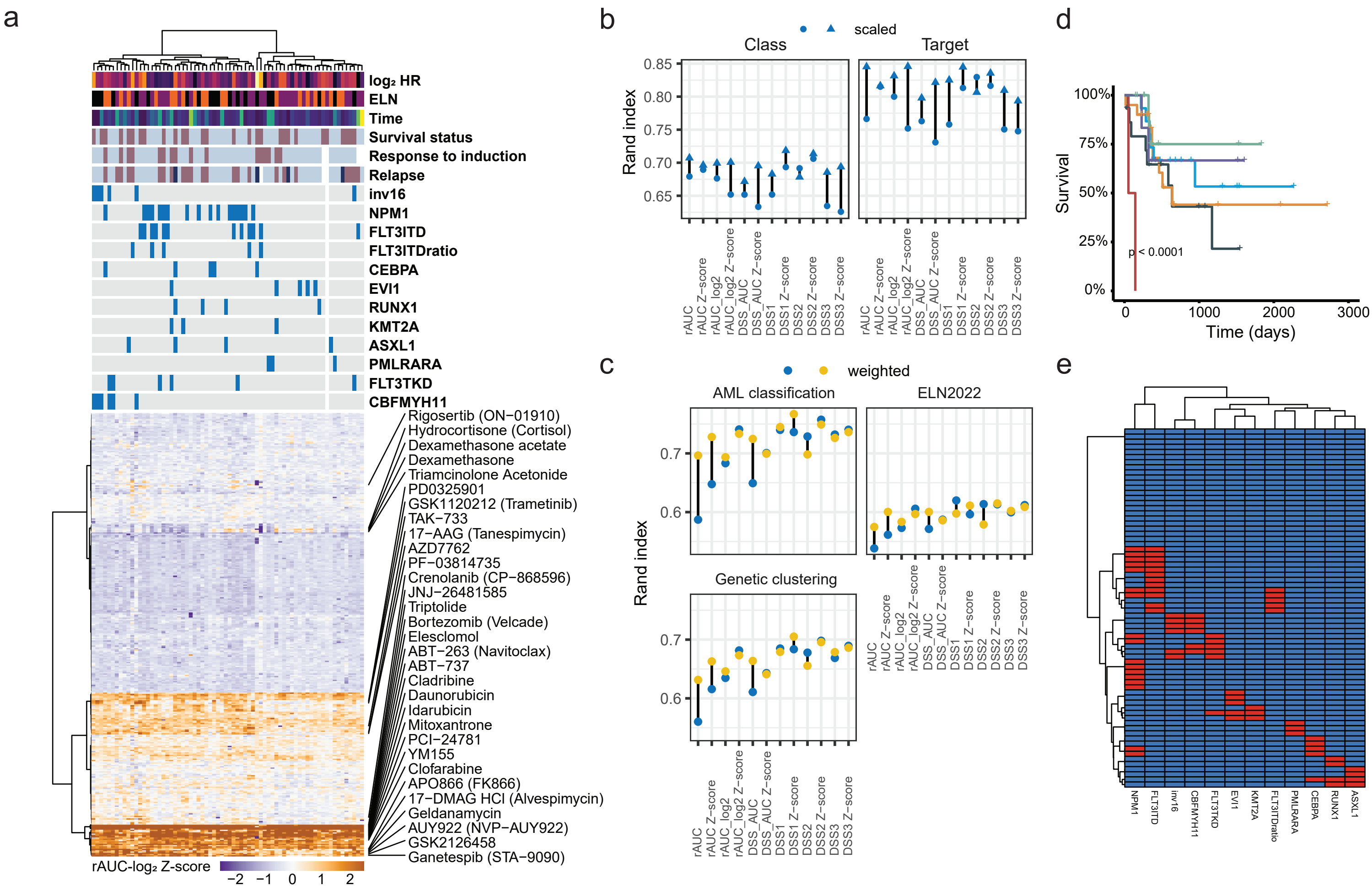
