## Supplementary material for "Clinical Forecasting using *Ex Vivo* Drug Sensitivity Profiling of Acute Myeloid Leukemia": Suppl. Methods

**Supplemental Methods**

*Ex vivo drug screening*

Blast cells were purified by LymphoprepTMgradient centrifugation (Stemcell) and cultured in Mononuclear Cell Medium (PromoCell C-28030) supplemented with 1% Penicillin + Streptomycin (PS) (Gibco,15140-122). A total of 10,000 cells (25 μL) was added to each well in pre-drugged 384-well plates (Greiner Bio-One) using a Multi Drop Combi peristaltic dispenser (Thermo Scientific). The number of cells was adjusted for specific samples with lower cell counts without a noticeable change in the outcome of the experiments. The Selleck Anti-Cancer Compound Library L3000 was used for drug screening, which consists of 349 anti-cancer drugs dissolved in dimethyl sulfoxide (DMSO) (see Suppl. Table S2 for the complete list). Compounds were pre-aliquoted in seven 384-well plates using five 10-fold dilution steps, reaching a concentration range from 1 nM to 10 μM. Eight positive (benzethonium chloride, BzCl) and negative (DMSO only) controls were added to each plate. The drug handling was done at the Biotechnology Center of Oslo using an Echo 550 (Labcyte Inc.). After incubation (72 hours at 37°C in a 5% CO2, humidified environment) relative cell viability was quantified using the CellTiter-Glo Luminescent viability assay (Promega) and an EnVision 2104 Multilabel plate reader (Perkin Elmer). The luminescence readout was measured in counts per second (CPS). Relative cell viability was computed from CPS values normalized to negative (DMSO) and positive (BzCl) controls within each plate.

*Dose-response analysis*

Computational analyses were performed in the statistical programming environment R (version 4.1.2). Relative cell viabilities (*y*) were computed from CPS values using min-max normalization to the median CPS of the positive and negative controls within each plate. All negative response values were adjusted to zero. Plate quality was assessed by a Z`-factor and differential z-score of the plate controls. The average drug sensitivity was measured as a normalized AUC of the raw relative viability response for every patient-drug combination using the formula:

$$rAUC=\sum_{c=2}^{5} \frac{y_{c}\left( logx_{c}-logx_{c-1} \right)}{logx_{5}-logx_{1}}$$

Here *x_c_* indicates a given concentration with relative viability *y_c_*, ranging from the lowest concentration *x_1_* to the highest concentration *x_5_*. The rAUCs were also negative log_2_-transformed to counter the skewness of the relative viability scale.

To score drug sensitivities using metrics based on dose response curve-fitting we used routines adapted from the Breeze application^22,25^. Briefly, Breeze transforms the scale to percent inhibition and the following parametrization of the Hill equation is fitted to the data:

$$f\left( x \right)=R_{min}+\frac{\left( R_{max}-R_{min} \right)}{1-{10}^{n\left( logEC50-logx \right)}}$$

For this only growth inhibitory responses are considered. All estimated EC50s are then adjusted to a value between the maximum and minimum concentration of the experiment. TEC50 is computed by setting EC50 to the maximum concentration for the models with R_max_ under 25%. Before further use, both EC50 and TEC50 were log-transformed and mean-subtracted per drug. The drug sensitivity score (DSS) is computed from an analytical solution to the integral$,$

$I=\int_{x_{t}}^{x_{5}} f\left( x \right)dx$,

where *x_t_* is the concentration at an activity threshold *t* set to 10%. Thus, DSS can only take positive values with $DSS1=\left( I-t\left( logx_{5}-logx_{t} \right) \right)/\left( \left( 100-t \right)\left( logx_{5}-logx_{1} \right) \right)$, $DSS2=DSS1/{log}R_{max}$, and $DSS3=DSS2\left( x_{5}-x_{t} \right)/\left( x_{5}-x_{1} \right)$. We also used Breeze to compute an AUC based on a LOESS curve-fit (referred to as DSS-AUC), which models both growth promoting and inhibitory responses.

Patient-wise standardization of drug sensitivity metrics (*s*), was performed followingly to compute a drug sensitivity z-score:

$z_{dr,pt}=\frac{s_{dr,pt}-\mu_{pt}}{\sigma_{pt}}$.

Here μ_pt_ and σ_pt_ is the drug sensitivity mean and standard deviation per patient respectively. For scaling of the drugs, a drug-wise standardization procedure was done over the drug sensitivity metrics or drug sensitivity z-scores. For analysis of differential drug sensitivity between relapsed and treatment-naïve samples, the difference in drug sensitivity z-scores was computed and statistical significance was evaluated using a paired t-test.

The similarities in drug sensitivity profiles between drugs or between samples/patients were measured using the Pearson correlation coefficient (PCC).

*Analysis of data confounders using PCA*

SVD was performed on scaled matrices of drug sensitivity metrics (or z-scores) such that $\left( s_{dr,pt}-\mu_{dr} \right)/{\sigma_{dr}}=\sum_{k} u_{dr,k}\sqrt{\lambda}_{k}v_{k,pt}$, where *u_dr,k_* and *v_k,pt_* are components of ***u****_k_* and ***v****_k_* which are left and right singular vectors of the principal axis *k* with variance $\lambda_{k}$. The variance explained by each principal component was computed as $\rho_{k}={\lambda_{k}}/{\sum_{k} \lambda_{k}}$. The association of specific sample characteristics with a principal component was assessed using linear regression against ***v****_k_* and measured with the adjusted r-squared (${r^{2}}_{adj}$). The patient sample characteristics with the following features were evaluated: average CPS signal for DMSO and BzCl; average CPS noise measured as the coefficient of variation for DMSO and BzCl as well as their differential z-score; the number of curve-fit non-responders measured as the number of DSS3 values equal zero or EC50 or TEC50 values at maximum; curve-fit error features, which included the number of low-confidence curve-fits reported by Breeze, and the average log-EC50 standard error, mean absolute error and max residual error per patient; batch covariates, which included time period clusters for screen executions, protocol batches and blast source (PBMC or BM); biological/clinical covariates, which included genetic and cytogenetic features, age, sex, WHO patient performance status, ELN2022 risk stratifications, cliincal history (primary AML or secondary after antecedent myelodysplastic synedromes or other causes), and AML FAB classifications. Cumulative variance explained by a sample characteristic up to a component K was calculated as $\sum_{k=1}^{K} \rho_{k}{r^{2}}_{adj,k}$.

PCA on dose-response data was done such that $\left( y_{\left( dr,c \right),pt}-\mu_{\left( dr,c \right)} \right)/{\sigma_{\left( dr,c \right)}}=\sum_{k} u_{\left( dr,c \right),k}\sqrt{\lambda}_{k}v_{k,pt}$, where *y_(dr,c),pt_* represent the raw, processed or Hill fitted drug response or curve-fit residuals of a patient (columns) and a drug at a specific concentration (rows). When indicated, standardization was applied over the columns.

*RCPC*

To remove potentially confounding principal components, the drug sensitivity metrics (or z-scores) were reconstructed as ${s'}_{dr,pt}=\sigma_{dr}\sum_{k>l} u_{dr,k}\sqrt{\lambda}_{k}v_{k,pt}+\mu_{dr}$, where *l* represents the minimum number of principal components removed. RCPC testing was performed for different Cox models and different dataset sizes based on the feature pre-selection described earlier (Fig. 2E-F and S4A-D) and for Lasso trained on 50 randomly sampled datasets (Fig. 2G and S4E-H).
